# Maternally transferred monoclonal antibodies protect neonatal mice from herpes simplex virus-induced mortality and morbidity

**DOI:** 10.1101/2022.01.12.476098

**Authors:** Iara M. Backes, Brook K. Byrd, Chaya D. Patel, Sean A. Taylor, Callaghan R. Garland, Scott W. MacDonald, Alejandro B. Balazs, Scott C. Davis, Margaret E. Ackerman, David A. Leib

**Affiliations:** Department of Microbiology and Immunology, Geisel School of Medicine at Dartmouth, Lebanon, NH 03756, USA; Thayer School of Engineering, Dartmouth College, Hanover, NH 03755, USA; Ragon Institute of MGH, MIT and Harvard, Cambridge, MA 02139, USA

## Abstract

Neonatal herpes simplex virus (HSV) infections often result in significant mortality and neurological morbidity despite antiviral drug therapy. Maternally-transferred HSV-specific antibodies reduce the risk of clinically-overt neonatal HSV (nHSV), but this observation has not been translationally applied. Using a neonatal mouse model, we tested the hypothesis that passive transfer of HSV-specific human monoclonal antibodies (mAbs) can prevent mortality and morbidity associated with nHSV. The mAbs were expressed *in vivo* by vectored immunoprophylaxis, or administered *in vivo* following recombinant expression *in vitro*. Through these maternally-derived routes or through direct administration to pups, diverse mAbs to HSV glycoprotein D protected against neonatal HSV-1 and HSV-2 infection. Using *in vivo* bioluminescent imaging, both pre- and post-exposure mAb treatment significantly reduced viral load. Administration of mAb also reduced nHSV-induced behavioral morbidity, as measured by anxiety-like behavior. Together these studies support the notion that HSV-specific mAb-based therapies may prevent or improve HSV infection outcomes in neonates.

**Graphical Abstract:** 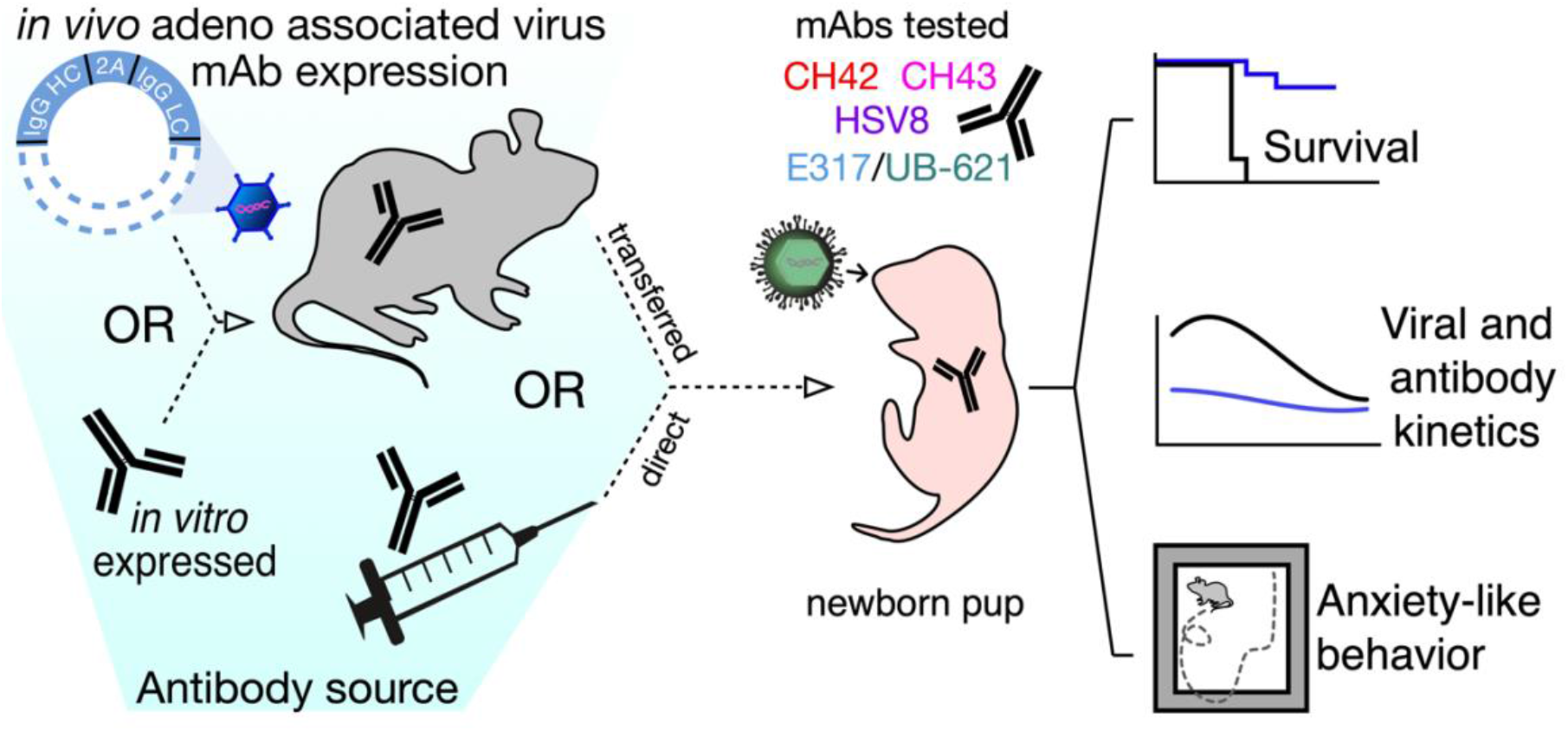

Different antibody sources were used to maternally-transfer or directly administer HSV-specific mAbs to mouse pups. Neonatal mice were challenged with wild type or bioluminescent virus before or after mAb acquisition. Following infection, pups were assessed for survival, virus-induced bioluminescence and anxiety-like behavior as a measure of neurological morbidity. Efficacy was time and mAb dependent. Notably, all HSV-specific mAbs prevented nHSV-associated mortality.

## Introduction

Among perinatal viral infections, neonatal herpes simplex virus (nHSV) infection has the highest infant mortality rate (1, 2). Transmission most often occurs during birth or via close contact with an infected individual in the first few days of life (3). It is estimated that there are 14,000 cases of nHSV per year globally (4) and recent studies suggest rising incidence in the United States (5). nHSV presents clinically in three forms: skin, eye, and mouth (SEM), central nervous system (CNS), and disseminated disease in the visceral organs. Despite aggressive antiviral treatment with acyclovir, mortality subsequent to disseminated disease remains high (30%), and CNS disease is associated with ~70% neurological morbidity and 25% mortality (6, 3, 7). This disease burden highlights the urgent need to augment clinical interventions in nHSV infections. Promisingly, maternal infection and serostatus is highly predictive of nHSV acquisition. The greatest risk (30-50%) is to vaginally delivered neonates born to seronegative mothers with primary or first-time genital infection (3, 7–9). In contrast, only ~3% of infections result from reactivated genital HSV disease, suggesting that maternal seroconversion and subsequent placental and/or milk transfer of antibodies protect the neonate from HSV infection (7–10). While many factors inform the relative risk of infant infection (10), there is a strong association between high titers of functionally potent anti-HSV maternal antibodies (Abs) and the absence of disseminated infection in neonates (11). Supporting these clinical observations, maternal seropositivity induced by vaccination, infection, or passive transfer of polyclonal HSV-specific IgG during pregnancy, reduced the risk of mortality and/or neurological morbidity in mouse pups (12–14).

However, only certain HSV-specific mAbs targeting envelope glycoproteins prevent ocular (15), genital (16, 17), and systemic HSV disease in adult mouse models (18, 19). Given the high susceptibility of neonatal mice to HSV infection, it is of interest to investigate whether mAbs that show efficacy in adult mice also prevent disease in neonatal mice. In agreement with adult mouse studies, protection in newborn mice depends on epitope specificity, with protection afforded by select mAbs or polyclonal antigen-specific Ab preparations (20). Here we evaluated a set of four human IgG_1_ mAbs that target the viral entry mediator glycoprotein D (gD), which is found on the HSV-1 and HSV-2 envelope as well as the surface of infected cells (15, 21–23), for their efficacy in preventing HSV mortality and morbidity in a neonatal mouse infection model.

Given the evidence that polyclonal maternal antibodies are naturally protective against nHSV, we hypothesized that maternally transferred mAbs could have therapeutic utility. To address this, dams were given an injection of recombinant mAbs, or adeno-associated virus (AAV) vectored antibody delivery (24), a strategy that can protect adult animals from viral (25–27) and parasitic infections (28), but which has not been evaluated for its ability to provide transgenerational protection in the form of maternally transferred Abs. Together our work demonstrates that maternal or direct administration of mAbs can ameliorate or prevent this devastating neonatal disease, motivating clinical investigation.

## Results

### mAb UB-621 accumulates at the placental-fetal interface

While maternal Abs prevent nHSV mortality and morbidity (12, 29), their biodistribution in pregnant dams has not been fully elucidated. To preserve the complex anatomy of the placental-fetal interface we pursued hyperspectral imaging via whole body cryo-macrotome processing (30), which causes minimal disruption to these tissues (Figure 1). We administered fluorescently labeled UB-621 mAb to pregnant C57BL6 (B6) dams two days before sacrifice and tissue preparation. Two dams (naive and UB-621 infused) were immediately processed for cryo-imaging after sacrifice, while an additional dam was further dissected to prepare conceptuses for cryo-imaging. Robust fluorescent signal of tissues at the placental-fetal interface was detected in the cryo-imaged dam infused with fluorescent UB-621 as compared to the control dam that did not receive mAb (Figure 1A and 1B, Supplemental Video 1). To confirm that this signal originated from tissues at the maternal-fetal interface, conceptuses were individually harvested from the maternal uterus and individually imaged with different layers of placental-fetal and/or fetal tissues dissected (Figure 1C). These dissection experiments confirmed that mAb accumulated at the placental-fetal interface, and was also detected in fetal and maternal tissues (Supplemental Figure 1). Notably, high fluorescence was observed in a harvested conceptus where the visceral yolk sac, a tissue rich in expression of the neonatal Fc receptor (FcRn) necessary for IgG transfer (31), remained intact (Figure 1C, middle image). Overall, this technique allowed us to visualize IgG traversing the circulation of the pregnant dam and entering maternal and fetal membranes.

**Figure 1.**
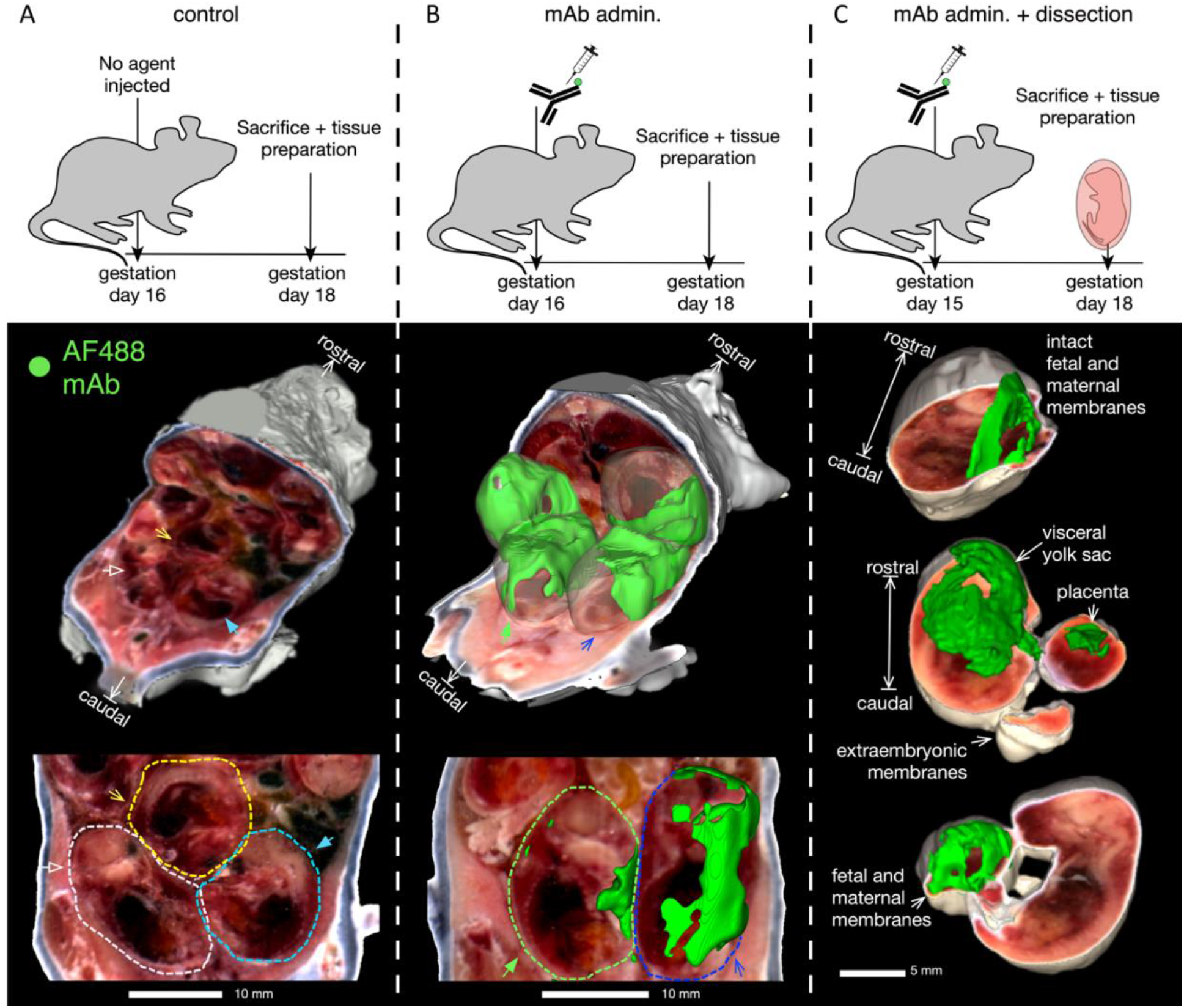
Fluorescently labeled mAb accumulates at the placental-fetal interface. To assess maternally administered Ab biodistribution, conjugated Ab was administered intravenously (IV) on day 15 or 16 of gestation, then 2-3 days later tissues were prepared for whole body imaging using the cryo-macrotome. **(A)** Background fluorescence levels in a pregnant dam not injected with conjugated Ab. **(B)** Accumulation of fluorescently labeled UB-621 Ab in a pregnant dam two days following IV administration. **(C)** Accumulation of fluorescently labeled Ab in conceptuses that were harvested from the murine uterus, with maternal and fetal layers removed as indicated.

### HSV mAbs targeting glycoprotein D protect neonatal mice from HSV-1 and HSV-2 mortality

The mAbs used in this study span the gD ectodomain, with epitopes close to the herpes virus entry mediator (HVEM) binding domain, and the Nectin (1 & 2) binding domains (Figure 2A). The gD:mAb interfaces between E317/UB-621, CH42 and CH43 have been resolved in detail through crystallography and alanine scanning (15, 21), while that of HSV8 is more broadly defined from binding experiments with truncated gD (32) (Figure 2B). Each of these mAbs protect from HSV infection in adult mouse models (Supplemental Table 1), and two (E317/UB-621 and HSV8) are currently in clinical trials (E317/UB-621: NCT02346760, NCT03595995, NCT04714060, NCT04979975, HSV8: NCT02579083). Like human neonates, mouse pups are highly susceptible to HSV infection, succumbing to infection at low viral doses relative to adult mice (33, 34). Therefore, we wished to determine if HSV gD-specific mAbs could protect mouse pups from HSV-1 infection. Pregnant dams were administered either CH42 or control IgG approximately 3-5 days before parturition, and pups were challenged intranasally with HSV-1 one day after birth (Figure 3A). Offspring of dams treated with CH42 showed significantly improved survival (p < 0.001) compared to offspring of control IgG-treated dams.

**Figure 2.**
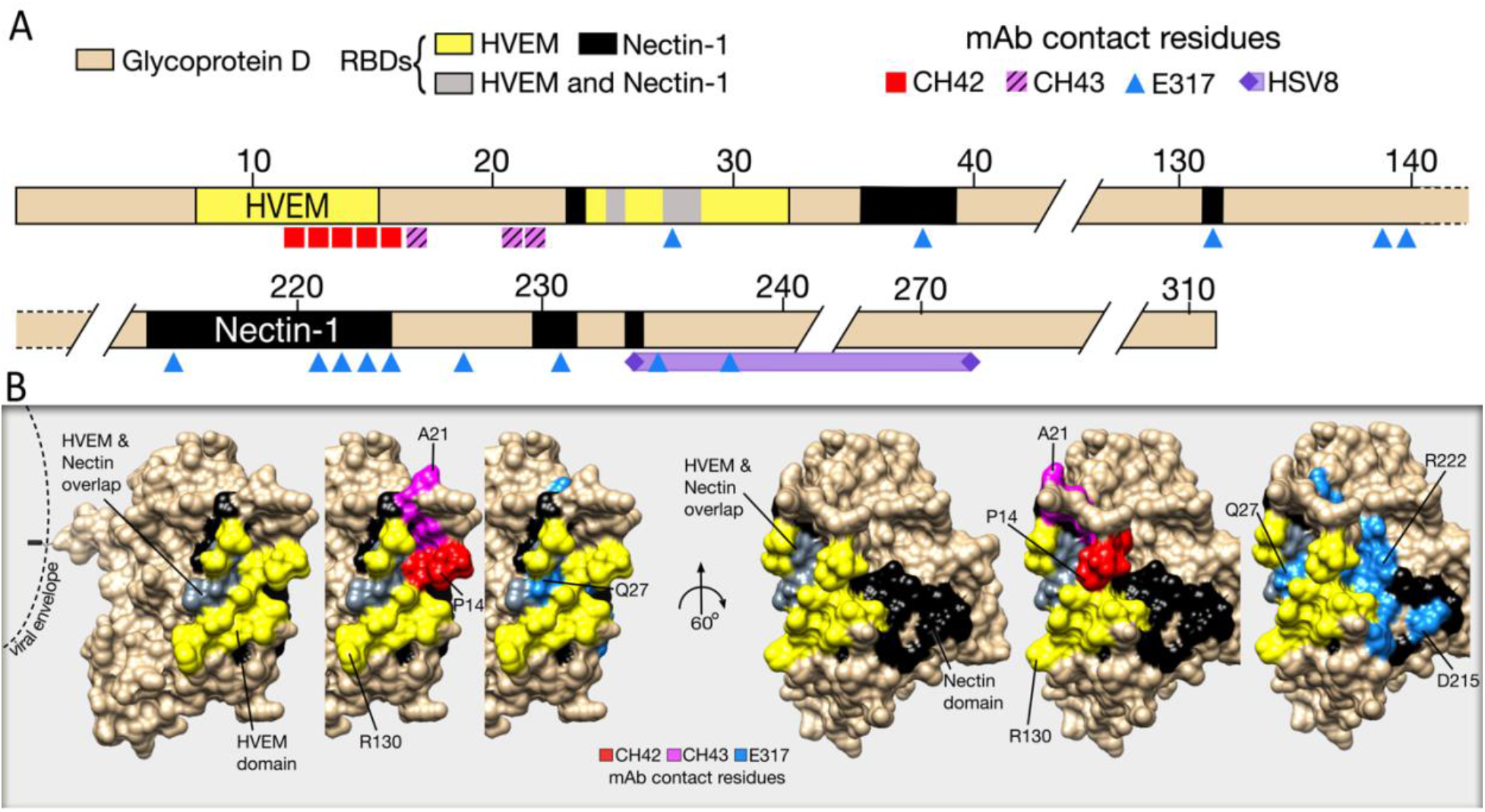
Epitope map of HSV-specific mAbs that target the viral entry mediator gD. (**A**) Linear representation of the gD extracellular domain with the HVEM binding domain (yellow) and Nectin-1 domain (black). UB-621 is derived from the original clone of E317. The exact contact residues of HSV8 are not known, but shown is an approximation based on reactivity against gD residues 235-275. (**B**) The gD ectodomain space filling structure (PDB: JMA1) with cell receptor binding domains and mAb epitopes denoted by specific colors shown in panel A, as follows: Red = CH42, pink = CH43, blue = E317, purple = HSV8.

**Figure 3.**
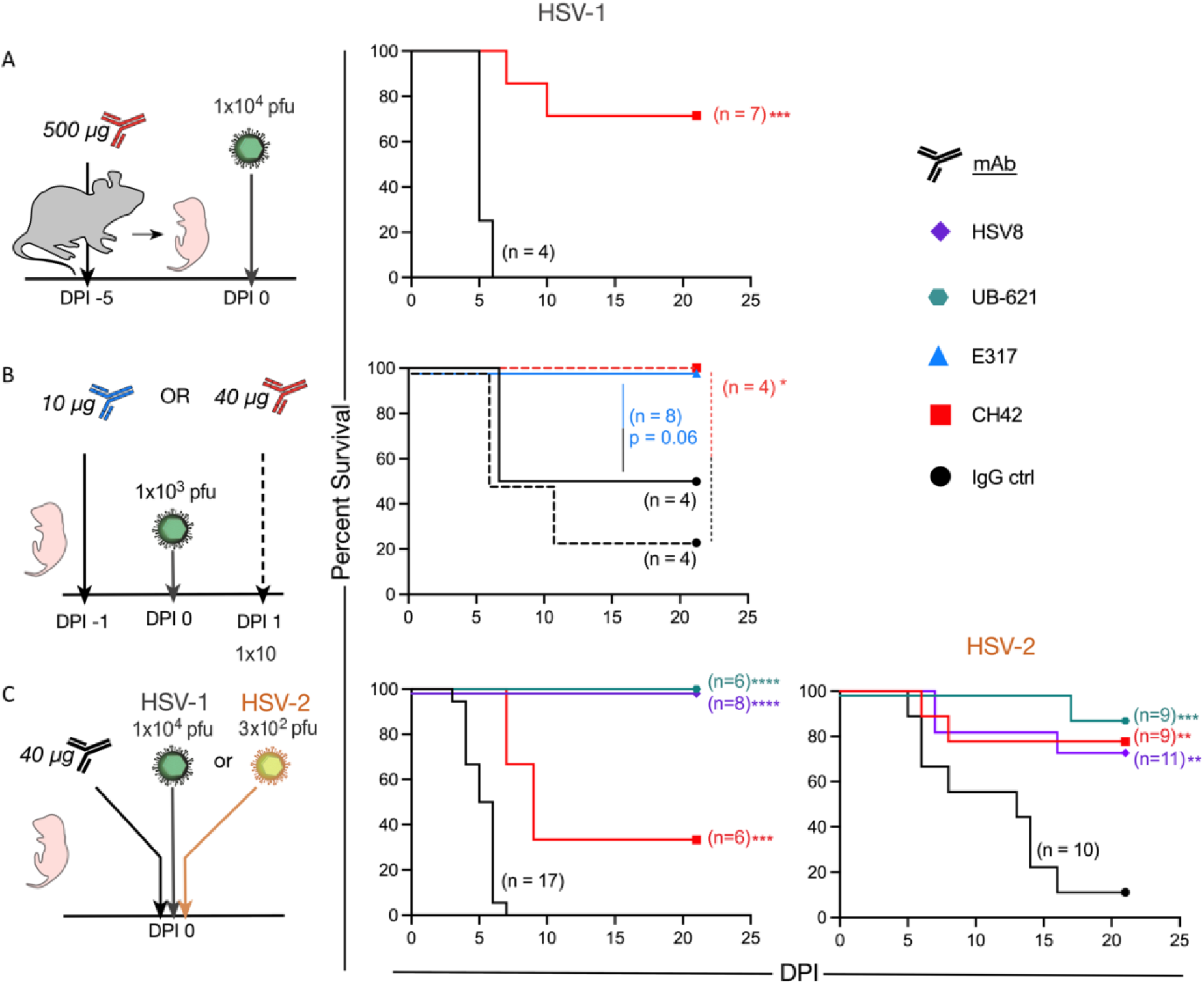
Prophylactic and therapeutic HSV-specific mAbs prevent nHSV-associated mortality. Abs were delivered intraperitonially to pregnant dams, or pups. Pups were challenged intranasally two days post-partum with indicated viral dose, then observed to DPI 21. (**A**) Survival of pups following CH42 (red) or IgG control (ctrl, black) administration to pregnant dams five days before infection. (**B**) Survival of pups following administration of E317 (blue) or IgG control (black) one day before infection in solid lines, or CH42 (red) or IgG control (black) one day post infection in dashed lines. (**C**) Survival of pups following HSV8 (purple), UB-621 (teal), CH42 (red), or IgG control (black) administration immediately before infection with HSV-1 or HSV-2. Statistical significance was determined by the Log-rank (Mantel-Cox) test; HSV-specific mAbs are compared to IgG control (ctrl). (* p < 0.05, ** p< 0.01, *** p<0.001). DPI, day post infection, PFU, plaque forming unit.

While prophylactic approaches for nHSV are desirable, we also sought to model therapeutic approaches to treat extant infections in neonates that may more closely model the clinical setting. To understand the prophylactic and therapeutic effect of mAb treatment, we administered E317 or control mAb one day *before* viral challenge, and given results with prophylactic maternal CH42 treatment, we administered CH42 or control mAb one day *after* viral challenge. Pups treated with E317 or CH42 exhibited improved survival (p=0.06 and p <0.05, respectively) relative to pups that received control IgG (Figure 3B).

We next assessed the protection afforded by mAbs currently being evaluated in clinical trials for adult genital HSV disease (HSV8 and UB-621) (22, 35) (E317/UB-621: NCT02346760, NCT03595995, NCT04714060, NCT04979975, HSV8: NCT02579083). HSV8, UB-621, CH42, or control IgG were administered to pups, then mice were immediately challenged with HSV-1. Both HSV8 and UB-621 mAbs completely protected pups from mortality following HSV-1 viral challenge (p < 0.001). In contrast, CH42 afforded partial protection compared to control IgG-treated pups (p < 0.01) (Figure 3C, left panel).

While HSV-1 genital disease predominates in the Americas and Western Pacific, and continues to rise as the etiologic agent of genital disease in high income countries, HSV-2 remains a significant cause of neonatal disease. Therefore, pups were treated with HSV8, UB-621 and CH42 as described above and challenged with HSV-2 (Figure 3C, right panel). Each HSV-specific mAb tested resulted in significantly improved (UB-621 p < 0.001, CH42 and HSV8 p < 0.01) survival of pups challenged with HSV-2 as compared to control IgG-treated pups. Collectively, these data showed that gD-specific mAbs can protect highly susceptible neonatal mice when administered via disparate routes, at different doses of antibody, and before and after viral challenge with either HSV-1 or HSV-2.

### mAb CH42 reduces CNS and disseminated viral replication

Disseminated disease results in the highest case fatality rate among nHSV clinical presentations, and despite aggressive antiviral treatment, has an unacceptably high mortality (30%) (7). We therefore assessed the impact of mAb therapy in the control of viral dissemination using bioluminescent imaging (BLI) to monitor viral replication and spread in real time (Figure 4). Dams or pups received CH42 mAb or control IgG either before or after challenge with recombinant HSV-1 that expresses luciferase (36, 37). We performed BLI daily from 2 to 8 days post infection. As expected (12, 29), virus was detected primarily in lungs, trigeminal ganglia, and brain (Figure 4, and Supplemental Figure 2). Among the different treatment and timing modalities tested, offspring of mAb-treated dams cleared the infections fastest compared to control IgG (p<0.001), with nearly undetectable bioluminescence even at the earliest timepoint assessed (Figure 4, A and D). When antibody and virus were administered simultaneously, or when antibody was administered one day following infection, a similar pattern was apparent: bioluminescence diminished more rapidly in CH42-rather than control IgG-treated pups over time and reached background levels by day 6 (Figure 4). While mAb prophylaxis of the pregnant dam resulted in the best viral suppression, co- and post-infection mAb treatment also conferred protection, demonstrating the efficacy of mAb therapy for nHSV.

**Figure 4.**
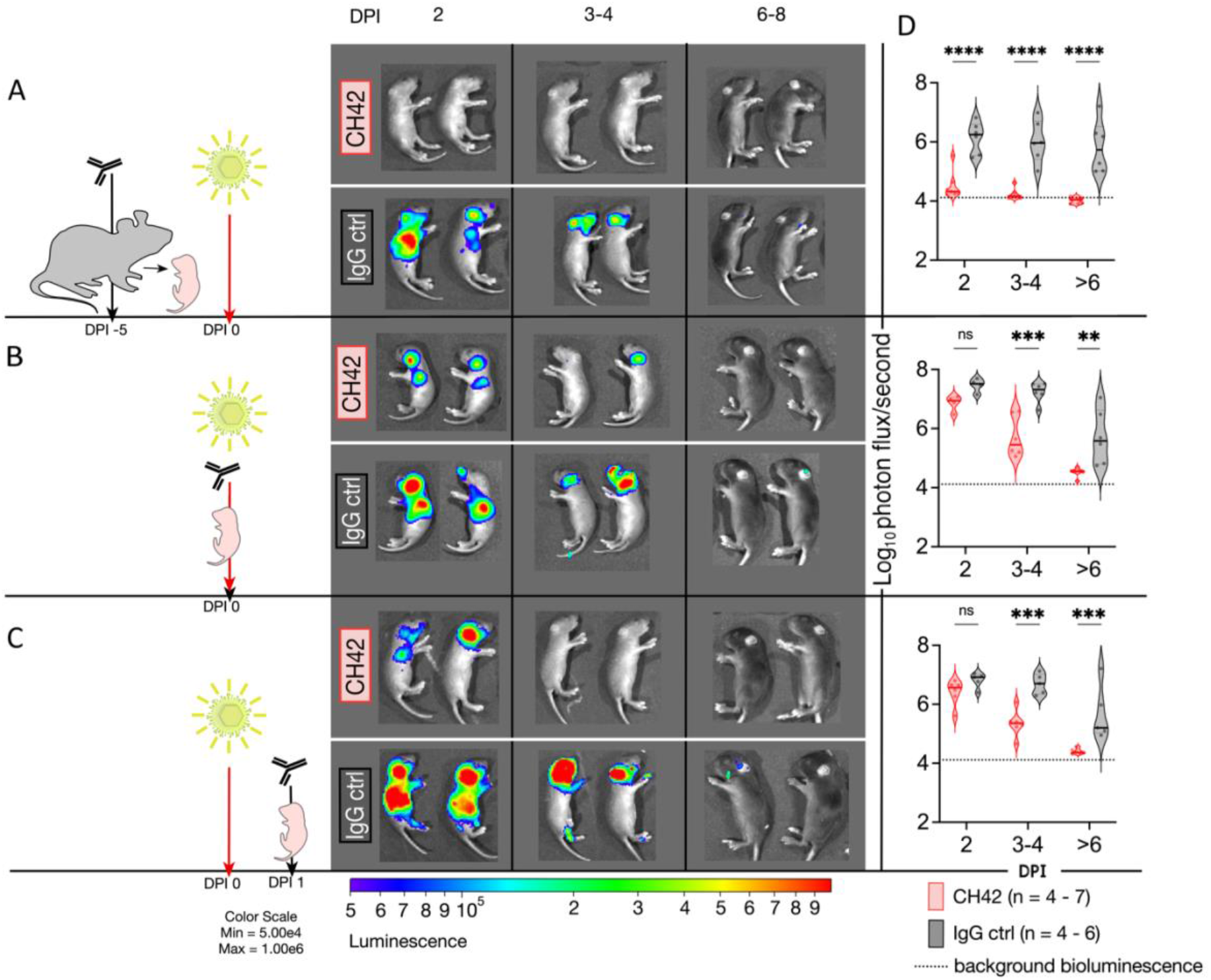
Administration of CH42 reduces viral dissemination. Pregnant dams or pups received injections of CH42 or IgG control (ctrl) antibody before or after viral challenge with a luciferase-expressing reporter HSV-1. Pups were imaged on DPI 2, then imaged until DPI 8. Representative images follow the same two pups sequentially. (**A**) Bioluminescence imaging of pups following CH42 or IgG control administration to pregnant dams five days before infection. (**B**) Bioluminescence imaging of pups following IP mAb administration and immediate subsequent viral challenge. (**C**) Bioluminescence imaging of pups following IP mAb administration one day after infection. (**D**) Quantification of the virally-derived bioluminescence shown in panels A, B & C. Statistical significance was determined by two-way ANOVA, and Šídák’s method for multiple comparisons (** p < 0.01, *** p< 0.001, **** p<0.0001).

### mAb immunotherapy reduces neurological morbidity in adult mice infected at birth

Neurological morbidity subsequent to HSV-1 infection of neonates was modeled using the Open Field Test (OFT, Figure 5A), which analyzes the innate exploratory behavior of mice as a measure of anxiety-like behavior (38). Mice are placed in an enclosed arena and the time spent in the periphery relative to the total exploration time (thigmotaxis ratio) is measured. Mice with anxiety-like behavior exhibit higher thigmotaxis ratios (39). Whereas scores of 0.5 are normal in B6 mice, HSV-infected neonates exhibit elevated thigmotaxis in adulthood (12, 14). We therefore tested whether HSV-specific mAbs could protect mice from exhibiting the anxiety-like behavior that follows neonatal infection. Offspring of control IgG-treated dams spent significantly (p < 0.05) more time in the periphery of the arena relative to progeny of CH42-treated dams (Figure 5B). Offspring of CH42-treated dams showed thigmotaxis scores close to 0.5 (Figure 5C), indicating that CH42 was able to protect mice from nHSV-induced behavioral morbidity.

**Figure 5.**
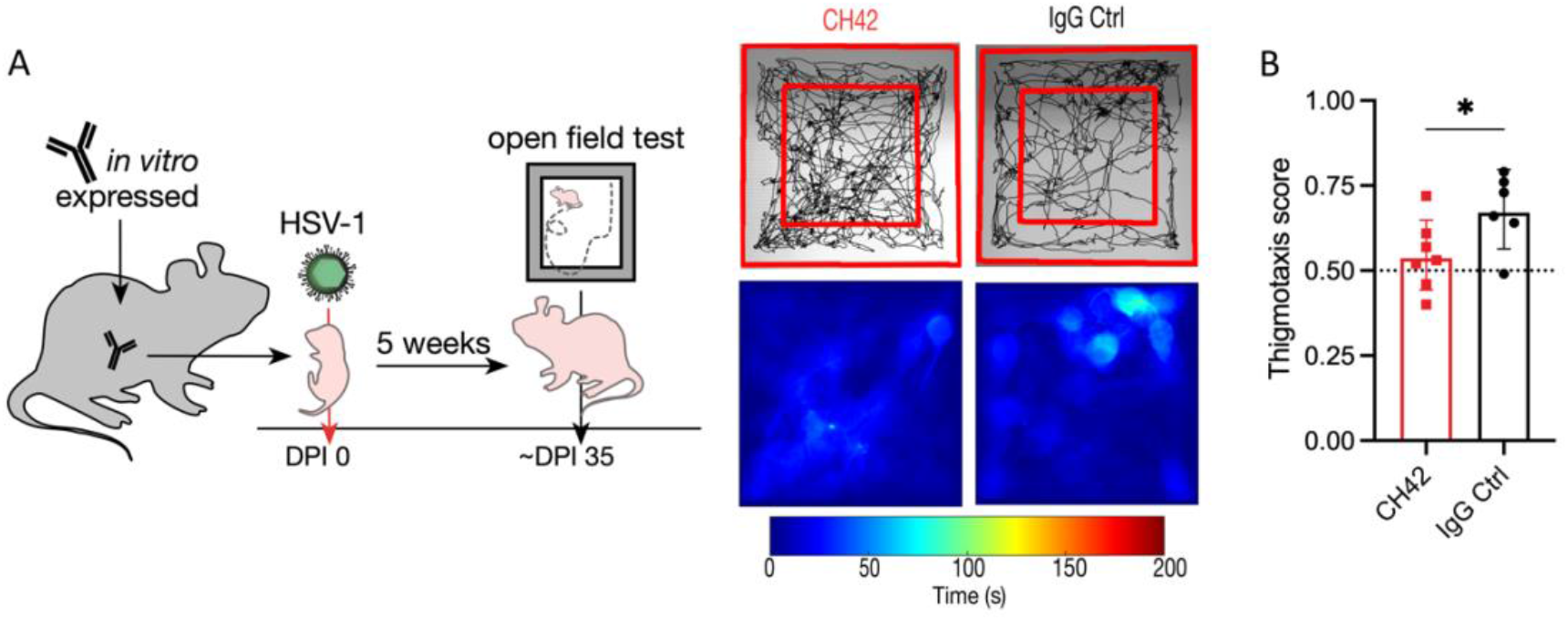
Offspring of HSV-specific mAb treated dams are protected from behavioral morbidity. Pregnant B6 mice were administered CH42 or IgG control (ctrl) mAbs, then progeny were challenged with luciferase expressing HSV-1, after 5 weeks mice were behaviorally assessed at 5 weeks. (**A**) Anxiety-like behavior analysis via the open field test (OFT) of adult mice infected on day two post-partum. Representative traces and heatmaps illustrate the pattern of movement, as well as the time spent in specific areas. (**B**) Thigmotaxis ratio of adult mice assessed for anxiety-like behavior via the OFT. Thigmotaxis is a measure of anxiety, a normal score = 0.5, higher scores indicate increased anxiety. Statistical significance was determined by unpaired t test (* p < 0.05).

### HSV-specific mAbs delivered via vectored immunoprophylaxis (VIP) provide trans-generational protection from nHSV mortality

Having shown that administration of mAbs to dams protects their pups from nHSV mortality, we sought to investigate vectored antibody delivery using AAV. Female mice received a single intramuscular injection of AAV encoding a human mAb (Figure 6A). Serum was obtained over a four-week period to confirm mAb expression (Figure 6B). Transduced dams expressed huIgG in the serum at different levels over a period of approximately 6 months and 3 pregnancies. Progeny of VIP-transduced dams had detectable levels of mAbs in serum (Figure 6C, Supplemental Figure 3A). Furthermore, mAb was effectively transferred throughout visceral organs and the nervous system (Figure 6C). Antibody biodistribution was similar to that observed in neonatal mice that received direct intraperitoneally (IP) injected mAb (Supplemental Figure 3B).

**Figure 6.**
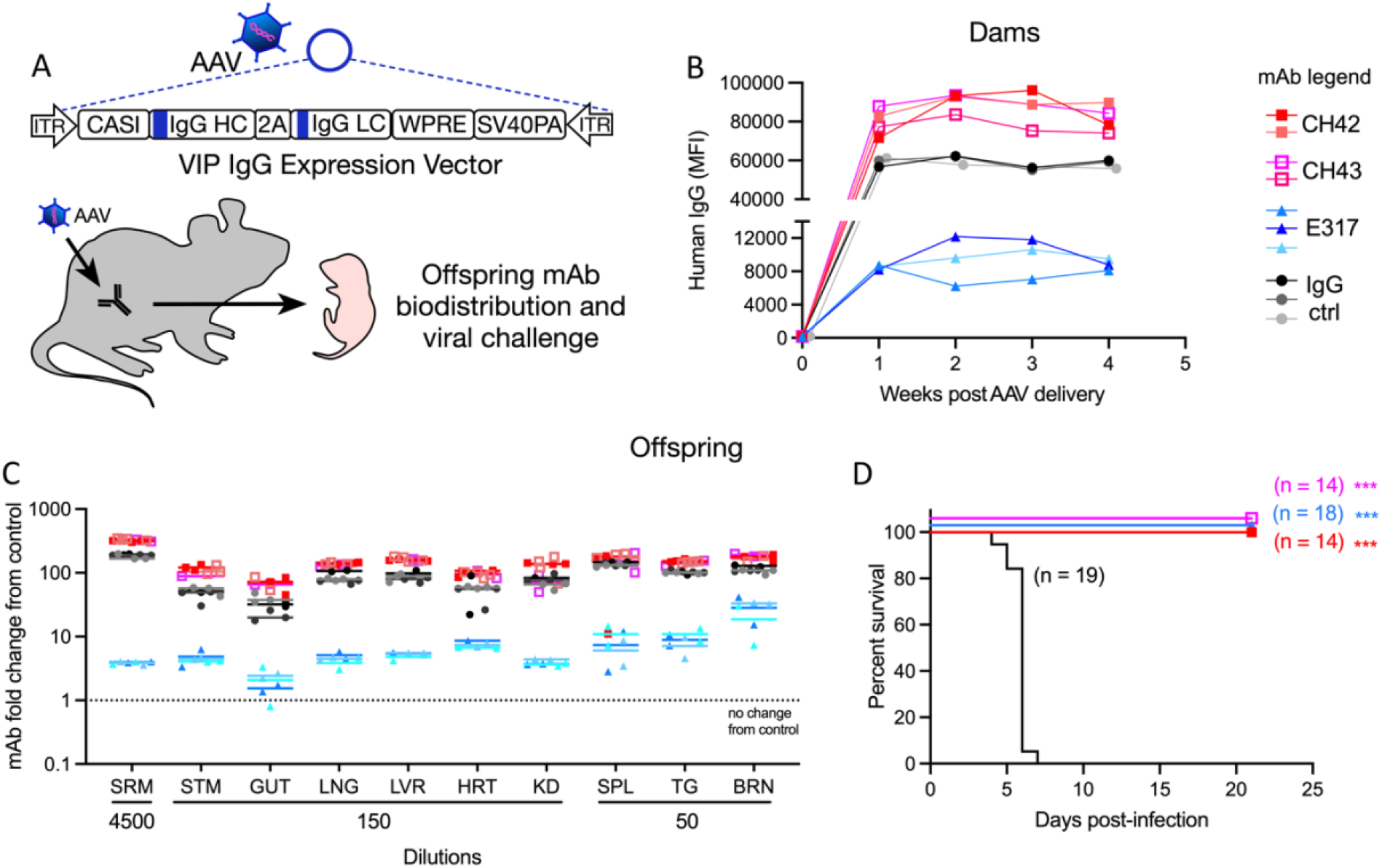
VIP-derived HSV-specific transferred mAbs protect pups from nHSV mortality. VIPs encoding 4 different mAb sequences were administered to female mice. Progeny were assessed for Ab transfer and protection from viral challenge. (**A**) Schematic of AAV human IgG expression vector structure and experimental approach. (**B**) Detection of *in vivo* expressed huIgG in the serum of VIP-administered female mice from week 0 through 4. (**C**) Biodistribution of human IgG in the viscera, brain, trigeminal ganglia and serum of offspring of VIP-treated dams. Signal is reported as the fold increase in human IgG in treated pups relative to untreated controls. (**D**) Survival of progeny of VIP-administered dams challenged with 1×10^4^ PFU of HSV-1 two days post-partum. Statistical significance was determined by the Log-rank (Mantel-Cox) test; HSV-specific mAbs are compared to IgG control (ctrl). (*** p< 0.001).

Finally, to assess whether the transferred mAbs were sufficient to protect pups from lethal viral challenge, progeny of VIP-transduced dams were challenged with HSV-1 two days postpartum, and monitored until weaning at three weeks of age. Despite variation in levels of transferred mAb, progeny of HSV mAb VIP-transduced dams were completely protected from mortality, while progeny of control IgG VIP-transduced dams rapidly succumbed to infection (Figure 6D). These findings demonstrate that antibodies produced *in vivo* in dams via VIP and subsequently transferred to pups were protective (*p* < 0.001). Moreover, although pups from E317 VIP-transduced dams received considerably less mAb, they were equally protected from disease as pups receiving 10-50-fold higher levels of mAbs from their mothers. Together, these findings further underscore the promise of diverse mAb delivery strategies in protection against nHSV-induced morbidity and mortality.

## Discussion

A great deal of evidence supports the hypothesis that Abs protect humans and mice against nHSV (9–13, 20, 29, 40–42). Several mAbs, some used in this study, ameliorate acute HSV infection in adult animal models (15–17, 20, 43). Despite this evidence, however, little progress has been made towards Ab-based therapies to treat nHSV in the clinic, and prior trials have relied upon acyclovir and its derivatives. Our experiments demonstrated that diverse mAbs targeting gD protect neonatal mice from lethal viral challenge and neuro-behavioral pathology. Among mAbs tested, HSV8 (35) and UB-621 (22) are currently being evaluated for prevention and/or reduction of adult HSV genital disease (clinical trial identifiers: NCT02579083, NCT04714060, NCT04165122). In light of our results, these mAbs should be considered for the prevention or amelioration of HSV in susceptible neonates.

Clinically, early therapeutic intervention is critical in the setting of nHSV, and delayed acyclovir treatment is associated with increased in-hospital death and morbidity (44). Likewise, our results underscore the importance of early intervention in the setting of mAb-based treatment. Among clinical presentations, disseminated disease has the highest mortality, and CNS disease the highest morbidity (6). Use of the intranasal route of infection, which often results in viral dissemination (12, 29, 40), allowed us to discern if mAbs could reduce viral load in multiple tissues. Maternal mAb administration, either by direct infusion of Ab or through VIP gave near-complete protection to pups from nHSV. These results are consistent with the idea that early prophylactic treatment provides optimum protection. Different mAbs afforded variable protection when directly administered to pups. These differences may be attributed to distinct mechanisms of Ab action (e.g., neutralization, antibody-dependent cellular cytotoxicity, or other effector functions), and/or differences in epitope affinity or specificity. More specifically, consistent with differences in survival between HVEM and Nectin knockout mice challenged with HSV (45–47), the differing receptor binding domains targeted by these mAbs could explain this variation. Overall, we conclude that the mAbs tested in this study are effective, and that the mode and timing of mAb administration are key determinants of protection. These data provide strong support to the idea that pre- and post-exposure mAb administration protects from the acute and long-term sequelae of nHSV infection.

Current clinical strategies to prevent nHSV involve administration of antivirals to mothers with a history of genital herpes or, in cases of primary HSV infection, cesarean section (10, 48). Importantly, our model uses maternal Ab therapy, which has the potential to protect both mother and neonate. Collectively, we show that mAbs transfer to neonates and distribute to visceral organs and the nervous system. This observation is consistent with the distribution of natural maternally-derived mouse and human Abs (14, 29). In agreement with the hypothesis that Abs protect neural tissues from viral spread (49–51), we see that maternal administration of mAb protected mice from neuro-behavioral pathology. These results represent an exciting opportunity to pursue maternal mAb-based interventions to prevent congenital and perinatal disease in at-risk populations. Adapting the AAV-vectored delivery of Ab, which is safe and durable in humans (52), could provide an avenue for maternal mAb transfer throughout gestation, potentially protecting both mother and baby. Hyperimmune globulin represents an alternative strategy to provide HSV-specific Abs to the maternal-infant dyad, which may also protect from nHSV infection. Recently, this approach was tried to prevent congenital cytomegalovirus (CMV) infection via maternal treatment with CMV hyperimmune globulin. Ultimately, this strategy did not prevent congenital CMV, and treatment with hyperimmune globulin raised concerns over adverse obstetrical outcomes. These adverse outcomes were not statistically significant, but they were consistent with previous studies that also detected slightly higher risk in preterm birth, preeclampsia, and reduced fetal growth (53, 54). In contrast, administration of mAbs during pregnancy has not been associated with these risks (55). Therefore, maternal mAb delivery may prove useful for other congenital and perinatal infections, especially for pathogens for which vaccination has failed or not been attempted.

Infectious disease accounts for approximately one-third of newborn deaths worldwide. Antenatal and perinatal infections of bacterial and viral etiology, including but not limited to Group B Streptococcus (GBS), CMV, and HSV have devastating consequences. These infections result in significant neonatal death and life-long disability of survivors (6, 7, 56–58). Encouragingly, widespread screening and antenatal antibiotic use have significantly reduced the incidence of neonatal GBS, demonstrating that infection in early life can be circumvented if good therapies exist (59). Viral infections such as CMV and HSV remain a significant danger to neonates. While antiviral treatments have significant benefits in reduction of mortality, they are unable to prevent long term sequelae of nHSV. In the absence of additional interventions such as vaccines or mAbs, consortium and working groups are unlikely to recommend screening, leaving a gap in identifying affected pregnancies/newborns and failing to prevent significant morbidity and mortality even when diagnosed. Therefore, there is a great need to identify effective new therapeutic interventions for antenatal and perinatal infections, as they have tremendous potential to save lives and improve long-term quality of life.

## Methods

### Mouse procedures

C57BL/6 (B6) and B cell insufficient muMT (B6.129S2-*Ighm^tm1Cgn^*/J) mice were purchased from The Jackson Laboratory (60). muMT mice were used in a subset of experiments to attribute protection to administered mAb, but results were interchangeable with the B6 mice which were therefore used for follow-up experiments. Blood collection was via cheek bleed from the mandibular vein with a 5mm lancet for weanlings and adults, or a 25 G needle for 1-2 wk old pups. Animals < 1 wk of age were euthanized prior to decapitation for blood collection. Blood samples were allowed to clot by stasis for ≥15 min. and then spun at 2000 × *g* for 10 min. at 4°C and supernatants collected and stored at −20°C. mAbs were administered IP to pups in 20 μL mAb were administered IP. to pregnant dams in volumes between 0.350 − 1 mL. For imaging studies, pups were injected IP with 20 μl of 15 mg/mL D-luciferin potassium salt, placed in isoflurane chamber, and moved into the IVIS Xenogen with a warmed stage and continuous isoflurane. Pups were typically imaged 2 days post-infection and serially imaged every other day to monitor bioluminescence. Endpoints for survival studies were defined as excessive morbidity (hunched, spasms, or paralysis) or >10% weight loss.

### Monoclonal antibodies

CH42 and CH43 plasmids were kindly provided by Dr. Tony Moody (Duke University). When expressed *in vitro*, CH42 contained the Fc mutation known as AAA (S298A/E333A/K334A), which enhances antibody dependent cellular cytotoxicity (61). E317 is the original clone of the clinical drug product UB-621; its heavy and light chain variable sequences were derived from published amino acid sequences (18) and synthesized in-house. The heavy and light chain variable sequences were ordered as “gBlocks” (Integrated DNA technologies) and cloned to an IgG1 heavy chain backbone, and a kappa light chain backbone. In-house expressed antibodies were made through co-transfection of heavy and light chain plasmids in Expi293 HEK cells (Thermo Fisher) according to the manufacturer’s instructions. Seven days after transfection, cultures were spun at 3000 × g for 30 minutes to pellet the cells, and supernatants were filtered (0.22 μm). IgG was affinity purified using a custom packed 5 mL protein A column with a retention time of 1 minute (ie. 5 mL/min) and eluted with 100 mM glycine pH 3, which was immediately neutralized with 1 M Tris buffer pH 8. Eluate was then concentrated to 2.5 mL for size exclusion chromatography on a HiPrep Sephacryl S-200 HR column using an AktaPure FPLC at a flow rate of 1 mL/min of sterile PBS. Fractions containing monomeric IgG were pooled and concentrated using spin columns (Amicon UFC903024) to approximately 2 mg/mL of protein and either used within a week or aliquoted and frozen at −80°C for later use. HSV8 mAb was kindly provided by ZabBio, and UB-621, a clinical grade antibody preparation with the E317 gene sequence expressed in hamster ovary cells, was kindly provided by United Biopharma.

### Viral challenge

The wild-type viral strains used in this study were HSV-1 17syn+(62), HSV-2 G (kindly provided by Dr. David Knipe) (63). The bioluminescent luciferase-expressing recombinant virus HSV-1 17syn+/Dlux was constructed as previously described (37). Viral stocks were prepared using Vero cells as previously described (64, 65). Newborn pups were infected intranasally on day 1 or 2 postpartum with indicated amounts of HSV in a volume of 5 μl under isoflurane anesthesia. Pups were then monitored for survival, imaging, or behavior studies once adulthood was reached as appropriate. For survival studies, pups were challenged with 1×10^3^ or 1×10^4^ plaque forming units (pfu) of HSV-1 (Strain 17), and 3×10^2^ pfu of HSV-2 (Strain G) as indicated. For imaging studies, pups were challenged with 1×10^5^ pfu of HSV-1 17syn+/Dlux.

### Whole body cryo-macrotome imaging procedures

Conjugation of mAb UB-621 was as previously described (66). Briefly, 5 mg of mAb in 100 μL of PBS was incubated with 10 μL of filter-sterilized 1M sodium bicarbonate and 1 μL of 10 mg/mL AF488 NHS-ester (Lumiprobe) for 1 hr. at room temperature and protected from light. Buffer exchange was carried out with Zeba spin columns (Thermo). Conjugation was confirmed through flow cytometry and spectrophotometer readings before animal experiments were performed. B6 dams were bred for timed pregnancies, and on day 11 of gestation chlorophyll-free diet (MP Biomedical) was initiated to reduce autofluorescence. On day 16 of gestation 5 mg AF488 labeled UB-621 was administered via tail-vein, and 2 days later animals were sacrificed and prepared for cryo-imaging by OCT (Tissue-Tek) flooding and subsequent freezing at −20 °C. The hyperspectral imaging whole body cryo-macrotome instrument has been described previously (67). Briefly, the system operates by automatically sectioning frozen specimens in a slice-and-image sequence, acquiring images of the specimen block after each section is removed. For this study, we acquired brightfield and AF488 fluorescence volumes of each animal at a resolution of 150 μm in the sectioning direction and ~100 μm in the imaging plane. Hyperspectral fluorescence images were spectrally unmixed using known fluorophore and tissue spectral bases to isolate the AF488 signal in animal tissue. The acquired image stacks were then combined in an open-source software platform (Slicer 4.11) to generate high-resolution three-dimensional volumes of the brightfield and fluorophore distribution throughout whole-body animal models.

### Adeno-associated virus (AAV) production and procedure

AAVs encoding the heavy and light chain sequences of CH42, CH43, and E317, and control IgG mAbs with the same human IgG1 backbone were produced as previously described (24). A single 40 μl injection of 1×10^11^ genome copies of AAV was administered into the gastrocnemius muscle of B6 mice as previously described (24). Blood samples were obtained by cheek bleed to verify antibody expression.

### Assessment of mAb expression and biodistribution

A magnetic bead-based assay (68) was used to measure antibody expression and biodistribution. Beads were conjugated to anti-human antigen-binding fragment (Fab) to capture mAbs of interest. Briefly, anti-human IgG F(ab’)2 fragment (Jackson Immune Research) was conjugated to fluorescent microspheres (MagPlex-C Microspheres, Luminex Corp.) at a ratio of 6.5 μg protein/100 μL microspheres. Samples were incubated with microspheres (500-750 beads/well) overnight at 4°C and washed in PBS with 1% BSA, 0.05% Tween-20, and 0.1% sodium azide. Anti-human IgG PE (Southern Biotech) was incubated at 0.65 μg/mL for 45 min in PBS-TBN. The microspheres were washed and resuspended in 90 μl of sheath fluid (Luminex) and read using a Bio-plex array reader (FlexMap 3D, OR MAGPIX). The median fluorescence intensity (MFI) of the PE signal was determined for each sample at indicated dilutions. For biodistribution assessment signal is reported as the fold increase in PE signal in treated pups relative to untreated controls.

### Behavioral tests and analysis

Animals were transferred to a dedicated behavior testing room at least one week before tests began. Environmental conditions, such as lighting, temperature, and noise levels were kept consistent. Behavioral tests and analysis were performed by independent, masked operators. The Open Field Test was performed as previously described (12). Briefly, 5- to 7-week-old B6 mice were placed in the open field arena (30 cm × 30 cm) and allowed to habituate for 10 mins before data collection took place for an additional 10 mins. The movement of animals was recorded (Canon Vixia HFM52) and videos were analyzed using open-source software (69).

### Statistical Analysis

Prism 8 GraphPad software was used for statistical tests unless otherwise described. For survival studies, HSV-specific mAbs were compared to isotype controls using the Log-rank Mantel-Cox test to determine *p* values. For imaging studies, groups and time points were compared to each other via two-way ANOVA, with Sidak’s test for multiple comparisons to determine *p* values.

### Study Approval

Procedures were performed in accordance with Dartmouth’s Center for Comparative Medicine and Research policies, and following approval by the institutional animal care and use committee.

## Acknowledgements

We gratefully appreciate Audra J. Charron for editing of the manuscript and thoughtful discussion. We acknowledge Alexey Khalenkov for construction of HSV-1 17syn+/Dlux, and appreciate Jennifer Fields for aid in animal studies. We also thank members of the Leib and Ackerman labs for materials and/or helpful discussion.

These studies were supported by NIH grants to DAL (R21 AI147714-01 to DAL and MEA, PO1 AI098681, and RO1 09083). A.B.B. is supported by the National Institutes for Drug Abuse (NIDA) Avenir New Innovator Award DP2DA040254, the MGH Transformative Scholars Program as well as funding from the Charles H. Hood Foundation. This independent research was supported by the Gilead Sciences Research Scholars Program in HIV.

## Authorship Contributions

I.M.B., S.C.D., A.B.B., M.E.A., D.A.L. designed and analyzed the research. I.M.B., B.K.B., C.D.P., S.A.T., C.R.G., S.W.M. performed experiments, and/or made materials, engaged in discussion, assisted with animal care and data analysis. I.M.B. drafted the manuscript and prepared figures, M.E.A., and D.A.L. edited the manuscript and figure legends, other authors contributed comments and edits.

