## Supplemental Figures and Legends for "Maternally transferred monoclonal antibodies protect neonatal mice from herpes simplex virus-induced mortality and morbidity"

**Supplemental Materials**

| mAb | Findings and summaries | Sources |
|---|---|---|
| CH42 & CH43 | Reduced mortality & the establishment of latency in corneal HSV-1 challenge of adult mice. | Wang, J Virol., 2017<br>US20140302062 |
| E317 | Reduced mortality in IP HSV-1 challenge of adult mice. | Lee, Acta. Cryst. D, 2013<br>WO2010087813A1<br>US8431118B2 |
| HSV8 | Reduced mortality in IP HSV-1 challenge of adult mice.<br>Vaginal antibody safety trial: Safety study of monoclonal antibodies to reduce the vaginal transmission of herpes simplex virus and human immunodeficiency virus. Phase I completed. | Sanna, Virology, 1996<br>Politch, PLoS med., 2021<br>NCT02579083 |
| UB-621* | Phase 1 clinical trial for SC administration proving safe and well tolerated in healthy volunteers.<br>Phase 2 clinical trials for the treatment of recurring genital HSV approved in the US and China.<br>*expressed in Chinese hamster ovarian cells from original clone E317 | NCT02346760<br>NCT03595995<br>NCT04714060<br>NCT04979975<br>Clementi, Drug Discov., 2016 |

**Supplemental Table 1.** Summary of the mAbs used in this study. IP, intraperitoneal. SC, subcutaneous.

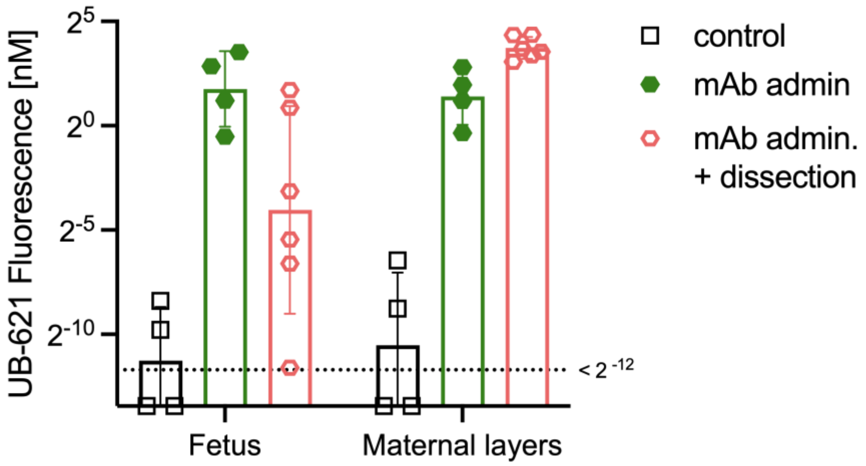


**Supplemental Figure 1. Fluorescently labeled mAb signals in fetal and maternal tissues measured by whole body hyperspectral imaging.** To assess maternally administered Ab biodistribution, conjugated Ab was administered IV on day 15 or 16 of gestation, then 2-3 days later tissues were prepared for imaging with the whole body cryo-macrotome. Control background fluorescence was determined in a pregnant dam not injected with conjugated Ab. Each dot represents the mean value of all voxels per slice segment for one animal, bar represents the geometric mean.

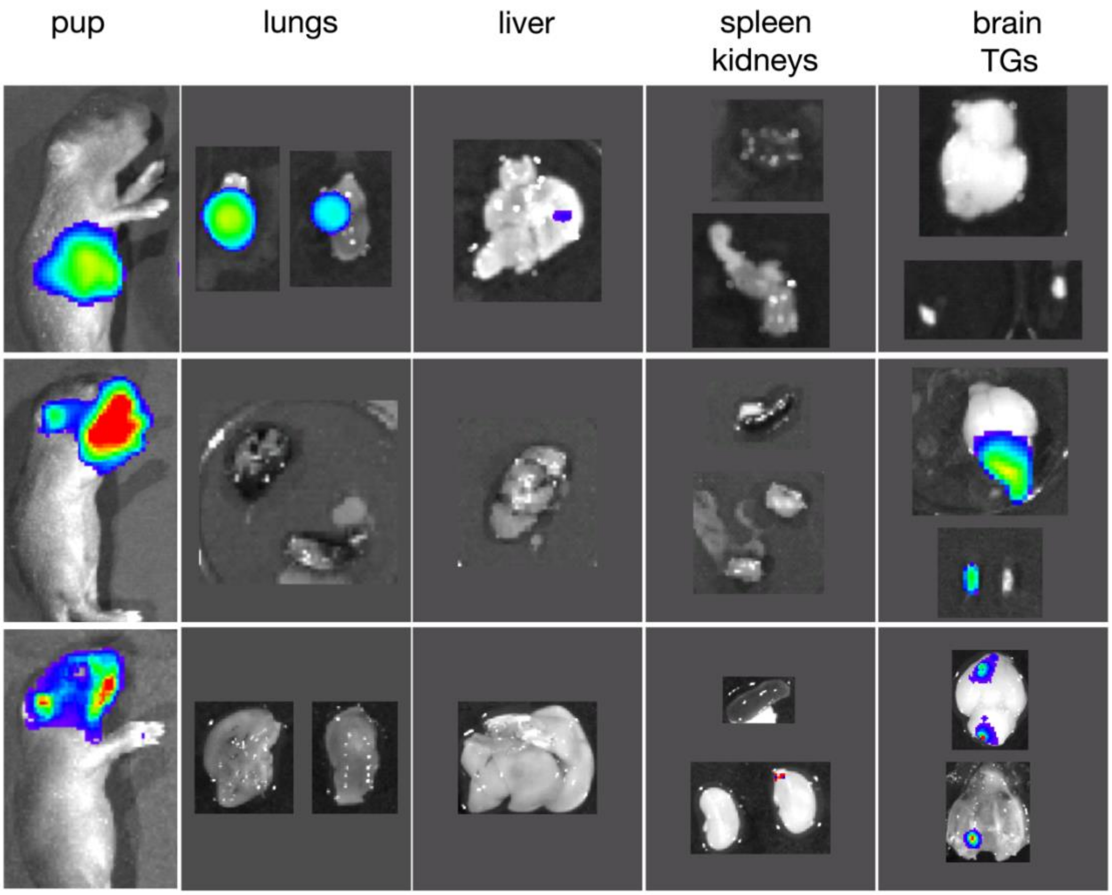


**Supplemental Figure 2 – Rostral bioluminescence derives from the brain and trigeminal ganglia.** Pups (n=3) were challenged with $1x10^5$ pfu of luciferase expressing HSV-1, then imaged following i.p. administration of 20 ul of luciferin (15 mg/mL). (**A**) Bioluminescence of pup imaged, and immediately sacrificed and dissected on DPI 2. (**B, C**) Bioluminescence of pups imaged, and immediately sacrificed and dissected on DPI 4. DPI= days post infection.

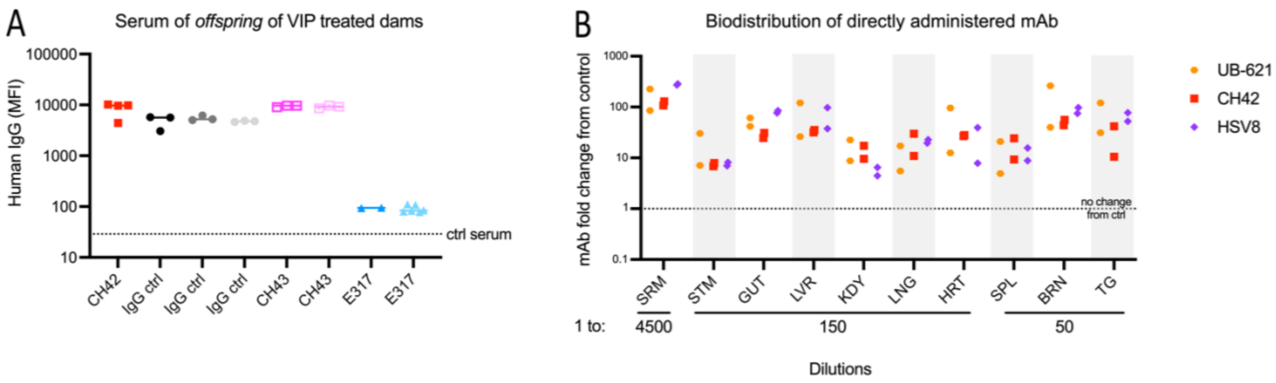


**Supplemental Figure 3 – Distribution of maternally transferred VIP-derived mAbs and directly administered mAbs in pups.** Custom bead-based assays were used to detect hIgG transfer or biodistribution**.** (**A**) Detection of *in vivo* expressed huIgG in the serum of 2-day old progeny of VIP-administered dams. Ctrl serum represents PE signal of the serum of progeny (n=3) of untreated naïve dams.  (**B**) Biodistribution of huIgG in the viscera, brain, trigeminal ganglia and serum of pups 48 hrs. after ip administration of mAbs. Signal is reported as the fold increase in huIgG in treated pups relative to untreated controls.
