## Supplemental Video 1 Legend for "Maternally transferred monoclonal antibodies protect neonatal mice from herpes simplex virus-induced mortality and morbidity"

**Supplemental video 1. Fluorescently labeled mAb signal in fetal and maternal tissues via whole body hyperspectral imaging.** Experimental set up as described in supplemental figure 1. Left video shows a naïve control dam that did not receive fluorescent mAb, right video shows dam administered AF488-conjugated UB-621.
